# Statistical testing in gene transcriptomic-neuroimaging associations: an evaluation of methods that assess spatial and gene specificity

**DOI:** 10.1101/2021.02.22.432228

**Authors:** Yongbin Wei, Siemon C. de Lange, Rory Pijnenburg, Lianne H. Scholtens, Dirk Jan Ardesch, Kyoko Watanabe, Danielle Posthuma, Martijn P. van den Heuvel

**Author notes:** Corresponding Author Martijn P. van den Heuvel, Prof., De Boelelaan 1085, W&N B-651, 1081 HV, Amsterdam.

## Abstract

Multiscale integration of gene transcriptomic and neuroimaging data is becoming a widely used approach for exploring the molecular underpinnings of large-scale brain structure and function. Proper statistical evaluation of computed associations between imaging-based phenotypic and transcriptomic data is key in these explorations, in particular to establish whether observed associations exceed ‘chance level’ of random, non-specific effects. Recent approaches have shown the importance of spatial null models to test for *spatial specificity* of effects to avoid serious inflation of reported statistics. Here, we discuss the need for examination of the second category of specificity in transcriptomic-neuroimaging analyses, namely that of *gene specificity*, examined using null models built upon effects that occur from sets of random genes. Through simple examples of commonly performed transcriptomic-neuroimaging analyses, we show that providing additional gene specificity on observed transcriptomic-neuroimaging effects is of high importance to avoid non-specific (potentially false-positive) effects. Through simulations we further show that the rate of reported non-specific effects (i.e., effects that are generally observed and cannot be specifically linked to a gene-set of interest) can run as high as 60%, with only less than 5% of transcriptomic-neuroimaging associations observed through ordinary linear regression analyses showing spatial and gene specificity. We explain that using proper null models that test for both spatial specificity and gene specificity is warranted.

## Introduction

A fast-growing number of imaging-genetic studies point out associations between the spatial patterns of gene transcriptome data and macroscale imaging-derived brain phenotypes (Richiardi et al., 2015; Wang et al., 2015a; Krienen et al., 2016; Anderson et al., 2018; Burt et al., 2018; Fornito et al., 2019; van den Heuvel et al., 2019). Whole-brain transcriptome data, such as the extensive Allen Human Brain Atlas (AHBA) serves as an invaluable quantitative reference to assess such transcriptomic-neuroimaging associations (Hawrylycz et al., 2012). Examples of such associations include genes related to oxidative metabolism to display transcriptional profiles in a similar spatial pattern as the degree of inter-module long-distance functional connectivity (FC) (Vertes et al., 2016) and hub connectivity (van den Heuvel and Sporns, 2013), as well as genes enriched for neuronal and synaptic connectivity to show transcriptional profiles that capture the architecture of brain functional networks (Krienen et al., 2016; Romero-Garcia et al., 2018b). Such transcriptomic-neuroimaging studies can also provide important new insight into the molecular background of macro-scale disease mechanisms. Brain transcriptional profiles of risk genes have been associated to patterns of disorder-specific brain changes revealed by neuroimaging techniques in several mental health conditions, for example schizophrenia (Romme et al., 2017), autism spectrum disorder (Romero-Garcia et al., 2018a) and major depressive disorder (Anderson et al., 2020), among others. These explorations have opened a new window for examining how genetic variants and molecular changes associated with brain disorders may relate to changes in brain structure and function.

As recently pointed out (Alexander-Bloch et al., 2018; Arnatkevic̆iūtė et al., 2019; Burt et al., 2020; Fulcher et al., 2020; Markello and Misic, 2021), caution is however advised in the statistical evaluation of gene expression patterns and imaging-derived features. An important point of these studies is that the commonly used linear model assumes independent observations. This is however not the case for brain gene expression data, as expression levels of neighboring regions tend to be often strongly correlated. The above mentioned studies have thus proposed important statistical methods to take such spatial autocorrelations into account, including null models that generate surrogate brain maps based on the parameterized variogram model (Burt et al., 2020) or based on spatial permutations (Alexander-Bloch et al., 2018). Implementations of such null models remarkably reduce false-positive findings (Fulcher et al., 2020; Markello and Misic, 2021), showing the extent of “spatial specificity” for the observed transcriptomic-neuroimaging associations. However, as also pointed by (Markello and Misic, 2021), these spatial null models still suffer from inflated false-positive rates.

What is not assessed in above spatial null models is the importance of evaluating whether an observed correlating transcriptomic-neuroimaging pattern goes beyond effects that one can expect from taking any other gene or set of genes, i.e., the extent of “gene specificity”. To address this type of specificity, other recent studies have proposed null models by generating null distributions of effects that are derived from random genes selected from the pool of all ∼20,000 genes. This null model is however not ideal. As importantly mentioned (Fulcher et al., 2020), it fails to take into account the level of co-expression among the genes of interest in the null condition, which may still lead to an inflation of statistical effects. Such a statistical bias strikingly increases when investigating a set of biologically relevant genes (like a set of genes related to a brain trait or a brain disorder), which are generally much higher co-expressed than a set of random genes (Wei et al., 2019). Null models based on co-expressed and biologically relevant random genes are thus required.

Given these various options to perform, it remains unknown how these different null models may impact statistical tests of transcriptomic-neuroimaging associations, and to what extent examinations of spatial specificity and gene specificity compensate for each other. Here, we compare and evaluate commonly used options for statistical evaluation of transcriptomic-neuroimaging associations, including the (most) commonly applied linear regression, null models that maintain spatial relationships, and null models on the basis of effects that occur amongst random genes. In line with recent studies (Burt et al., 2018; Arnatkevic̆iūtė et al., 2019; Fulcher et al., 2020), we point out that controlling for spatial specificity is important to reduce false-positive findings, but we further show that this is not enough: We suggest that further examinations of gene specificity using proper null models that account for random gene effects is equally important in minimizing bias towards non-specific results in transcriptomic-neuroimaging studies. We provide a toolbox to easily perform null-model evaluations for transcriptomic-neuroimaging studies.

## Results

We demonstrate the use and importance of different null models by means of discussing the analysis strategy and results of three common examples, followed by simulations of effects. We assess how the different null models (i.e., null models that give spatial or gene specificity) serve to identify possibly inflated statistical effects. For each example, we hypothesize that the transcriptional profile of a gene/gene-set of interest (GOI) relates to a brain phenotype, like the spatial pattern of brain atrophy (McColgan et al., 2018) or brain disconnectivity (Romme et al., 2017). We test a hypothesized association between the GOI and the brain phenotypes by examining the correlation between the expression pattern of the GOI across the cortex and the pattern of brain features, e.g., atrophy across the cortex (Figure 1a). We then test the statistical relevance of these associations using different strategies: 1) by spatial null models that provide spatial specificity (Figure 1b) and *2)* by random-gene null models that assess gene specificity (Figure 1c). We show that statistical evaluations based on null models that maintain spatial relationships do not necessarily provide gene specificity, and vice versa.

**Figure 1.**
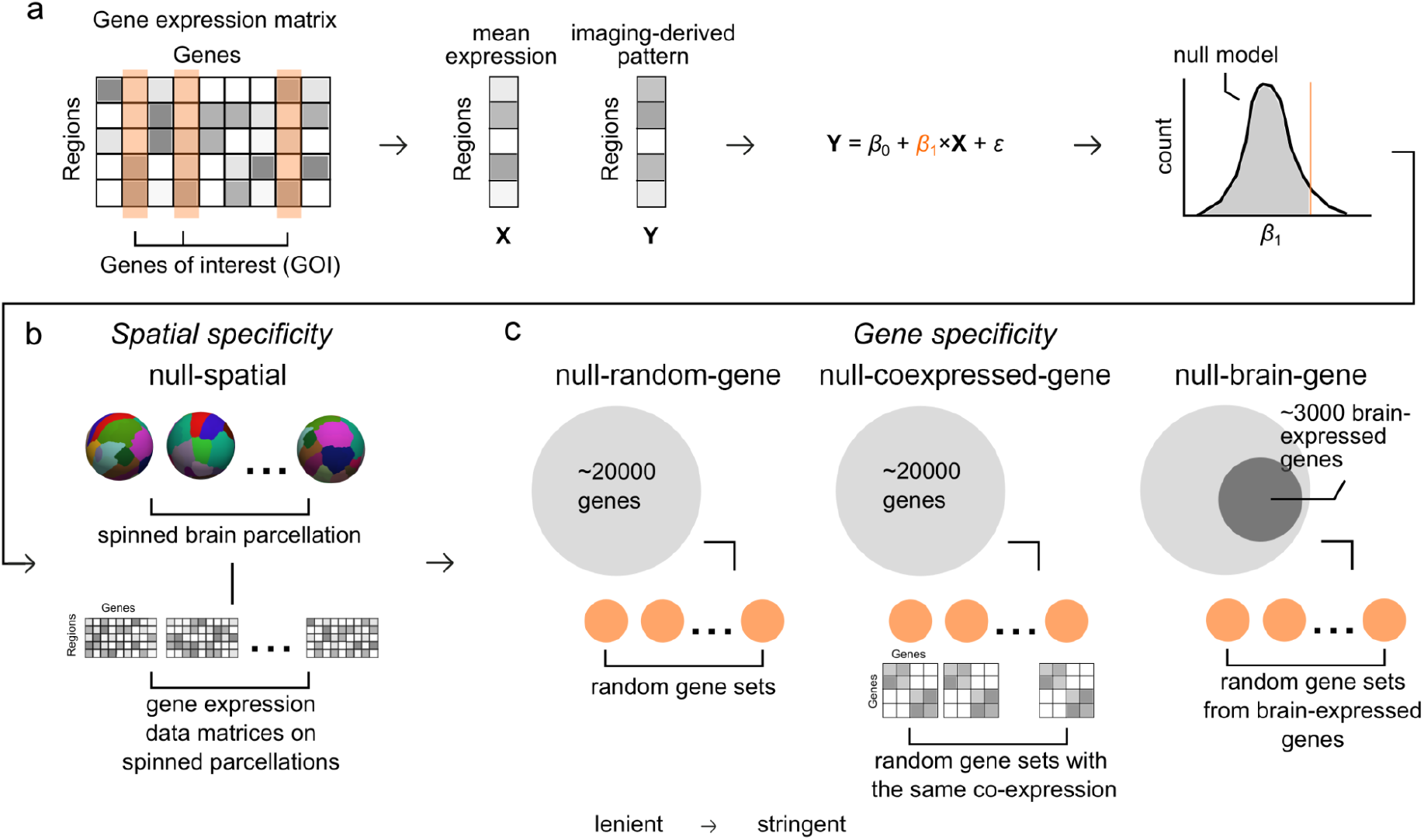
Approaches for statistical testing of overlapping patterns of gene transcription and imaging-derived brain phenotypes. (a) *Permutation test procedure*. The expression profile (**X**) of a user-defined set of the gene(s) of interest (GOI) is computed. The association between the expression profile and the imaging-derived pattern (**Y**) is assessed using linear regression. Permutation testing is used to examine whether the observed *β*_1_ is larger than null distributions of *β*_1_ derived from null models. Different statistical null models are possible: (b) *Examination of spatial specificity*. The “*null-spatial*” model (Alexander-Bloch et al., 2018) is proposed as a method to examine the spatial specificity of the observed associations. Randomized brain parcellations are obtained by spinning the inflated sphere of the real brain parcellation (1000 randomizations). A gene expression data matrix is then rebuilt using these randomized brain parcellations. The rebuilt gene expression data is used to re-evaluate transcriptomic-neuroimaging associations to generate null distributions. (c) *Examination of gene specificity*. Three null models (from more liberal to more stringent) are used to examine the gene specificity. *“null-random-gene”* model: random genes are selected from all ∼20,000 genes included in AHBA. *“null-coexpressed-gene”* model: random genes that conserve the mean co-expression level of the original genes are selected from all ∼20,000 genes included in AHBA. *“null-brain-gene”* model: random genes are selected from a subset of genes (2957 genes) that show up-regulated expression levels in brain tissues in contrast to other body sites.

### Example 1: genes related to cortico-cortical connectivity

We start by re-assessing a commonly examined relationship between the expression of genes important for neuronal connectivity and macroscale connectome organization (Krienen et al., 2016; Romero-Garcia et al., 2018b). We test this by focusing on 19 genes known to be enriched in supragranular layers of the human cerebral cortex [referred to as human supragranular enriched (HSE) genes], revealed by a previous study that compared gene expression profiles in different cortical layers between mice and humans (Zeng et al., 2012). These HSE genes are suggested to play a role in shaping long-range cortico-cortical connections of layer III pyramidal neurons and as such to show a spatial expression pattern that runs parallel to the organization of brain structural and functional networks (Krienen et al., 2016; Romero-Garcia et al., 2018b). The transcriptional profile of HSE genes is shown in Figure 2a.

**Figure 2.**
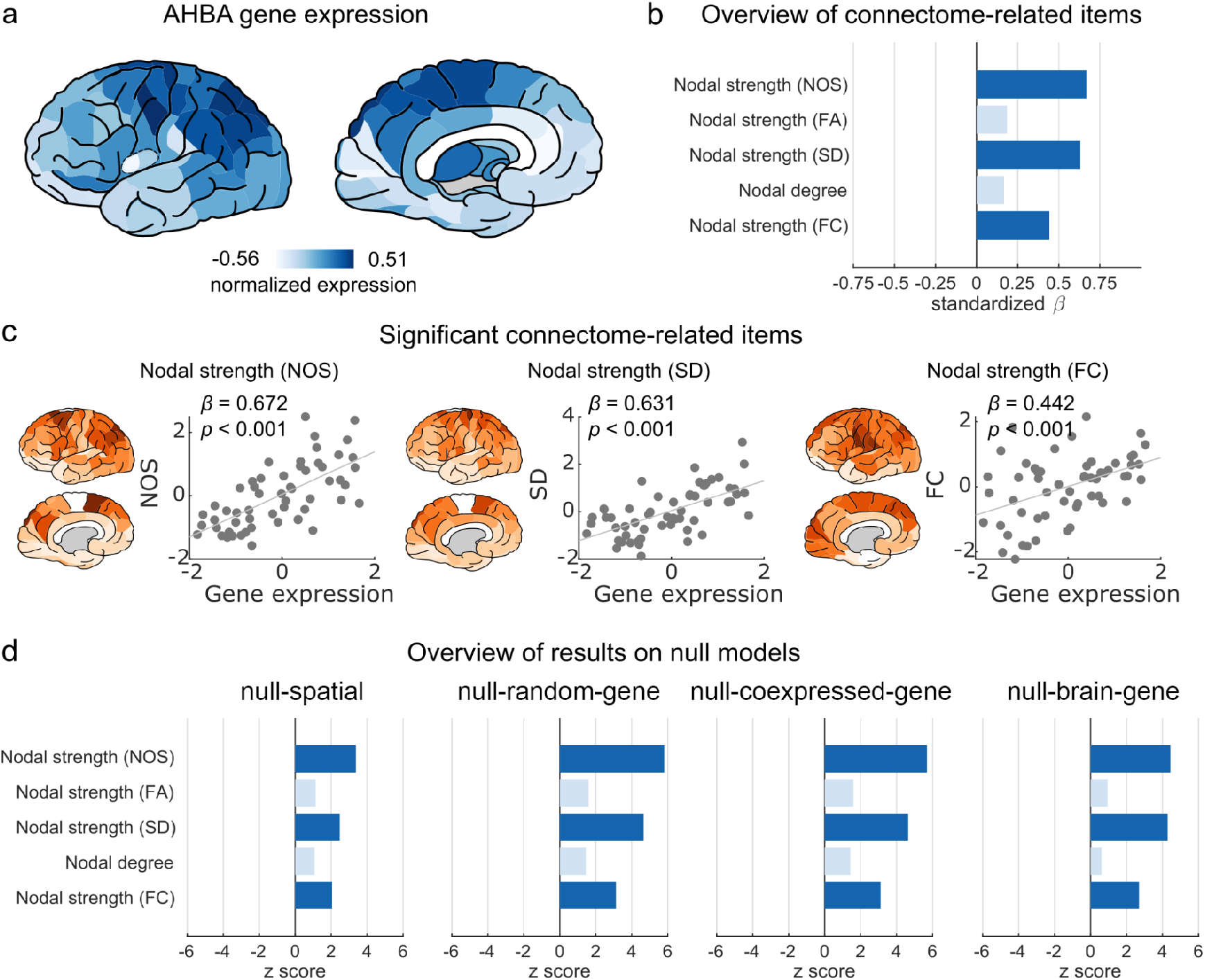
HSE gene expression and macroscale connectome properties. (a) Brain plots of normalized gene expression levels of HSE genes. (b) Overview of linear regression results between HSE gene expression profile and five imaging-derived connectome properties included in GAMBA. Dark blue indicates significant (*q* < 0.05, FDR corrected across 5 connectome traits). (c) Scatter plots for significant correlations between HSE gene expression and nodal strength of the structural (NOS-weighted, *β* = 0.672 and SD-weighted, *β* = 0.631) and functional connectome (*β* = 0.442). (d) Permutation testing results showing whether the observed effect size (*β* in panel b) is significantly beyond four distinct null distributions of effect sizes for null-spatial, null-random-gene, null-coexpressed-gene, and null-brain-gene models. Dark blue indicates *p* < 0.05 (two-tailed *z*-test).

#### Transcriptomic-neuroimaging overlap: linear regression

We first examine the spatial expression pattern of HSE genes in the context of macroscale brain connectome properties (Romero-Garcia et al., 2018b). Using simple linear regression we indeed note that the expression pattern of HSE genes is significantly associated with the pattern of connectivity strength of cortical areas – a measurement describing the extent to which a region is connected to the rest of the brain – under the assumption of ‘independent’ gene expressions in the brain [standardized beta (*β*) = 0.672, *p* < 0.001 for connectivity weighted by the number of streamlines (NOS); *β* = 0.631 *p* < 0.001 for connectivity weighted by streamline density (SD); false discovery rate (FDR) corrected for multiple testing across five connectome-related metrics; Figure 2b,c]. A similar association can be found between the pattern of HSE gene expression and the pattern of nodal strength of the functional connectome (FC; *β* = 0.442, *p* < 0.001; FDR corrected; Figure 2b,c), which is in line with the notion of a strong correspondence between structural and functional connectivity (Wang et al., 2015b).

#### Spatial specificity: null-spatial model

These findings thus confirm a potential association between HSE gene expression and macroscale connectome organization (Krienen et al., 2016; Romero-Garcia et al., 2018b), but it is worth checking whether such associations are *spatially* specific (i.e., not inflated by inter-regional auto-correlations of gene expression). As discussed recently (Alexander-Bloch et al., 2018; Arnatkevic̆iūtė et al., 2019; Burt et al., 2020; Fulcher et al., 2020), the commonly used linear regression method assumes independent observations, which means no correlation between values (i.e., expression levels) of the same variable [i.e., gene(s)] across different observations (i.e., regions). It is thus crucial to examine whether the observed associations are biased by potentially correlated expressions of neighboring brain regions, i.e., preserving spatial relationships across regions. To this end, an important new null model was introduced, namely the “null-spatial” model where transcriptomic samples are assigned to brain regions from randomized parcellations that are obtained by spinning the reconstructed sphere of the real brain parcellation (Alexander-Bloch et al., 2018), importantly preserving spatial relationships across brain regions (1,000 randomizations; see Methods, Figure 1b). The rebuilt gene expression data matrices are used to generate a null distribution of *β*, which is used to assess whether the original effect goes beyond the null condition. Using the null-spatial model, the associations between the expression pattern of HSE genes and the pattern of structural/functional connectivity strength are again found to be significant (NOS: *z* = 3.372, *p* < 0.001; SD: *z* = 2.473, *p* = 0.013; FC: *z* = 2.051, *p* = 0.040; FDR-corrected for multiple testing across 5 tests, Figure 2d). These effects thus significantly go beyond effects that one can expect using randomized brain regions with the same neighboring relationships, suggesting that the observed associations are specific to the spatial organization of brain regions.

#### Gene specificity

Spatial specificity however does not yet tell us anything about whether the observed effect is specific to this set of genes or might be present for all genes associated with the brain. We thus further want to examine whether the observed effects are also gene specific (i.e., not reflected by other genes in general). This is of particular importance as multiple examinations have pointed out general (global) posterior-anterior gradients of expression levels across brain areas (McColgan et al., 2018; Vogel et al., 2020), effects that may more reflect spatial patterns general to genes expressed in the brain, rather than reflecting patterns unique to the GOI. We thus argue that the next important step in the evaluation of the observed association(s) between transcriptomic profile(s) and spatial patterns of the imaging-derived brain phenotypes is to examine to what extent the computed linear correlations exceed effects that one could also observe when *any* other gene or set of genes would have been chosen, i.e., a set of not-of-interest genes.

#### Null-random-gene model

We employ permutations to generate null distributions of effect sizes, *β*, based on gene expression profiles of same-sized gene sets randomly selected from all genes (by default 10,000 permutations are used). We refer to this commonly used null model (Romme et al., 2017; Fulcher et al., 2020) as the “null-random-gene” model (Figure 1c). When the original *β* significantly exceeds this null model, it indicates the observed association cannot be found for all genes, but is unique for the GOI. For our HSE example, we show that the observed effect sizes for associations between HSE genes and nodal strength of structural and functional connectivity are significantly larger than effect sizes of random genes (NOS: *z* = 5.810, *p* < 0.001; SD: *z* = 4.660, *p* < 0.001; FC: *z* = 3.124, *p* = 0.003; Figure 2d).

#### Null-coexpressed-gene model

We (Wei et al., 2019) and others (Fulcher et al., 2020) have, however, further noted that when a set of genes (instead of a single gene) is examined, the co-expression level within the set of genes can influence findings resulting from the null-random-gene model. This is because the null-random-gene model underestimates the covariance of genes, something that is not conserved in a random mixture of genes (Fulcher et al., 2020). Therefore, we argue that it would be more informative to further include a null model for stricter statistical evaluation, comparing the observed effect size to the null distribution of effect sizes yielded by random genes conserving the same level of co-expression as the input GOI (referred to as “null-coexpressed-gene”; Figure 1c). For our HSE example, we use simulated annealing to look for random genes with the mean co-expression levels converging to those of HSE genes (see Methods). Application of this null model still shows significant associations between HSE gene expression and connectivity strength (NOS: *z* = 5.690, *p* < 0.001; SD: *z* = 4.640, *p* < 0.001; FC: *z* = 3.114, *p* = 0.002; Figure 2d).

#### Null-brain-gene model

We anticipate that the strongest usability of transcriptomic-neuroimaging examinations is to link brain transcriptomic data (like AHBA) to brain phenotypes (like MRI measurements). Most of such examinations will thus be centered on testing a GOI pre-selected based on the findings from previous brain related studies, e.g. GWAS of a disease phenotype like schizophrenia (Schizophrenia Working Group of the Psychiatric Genomics, Consortium, 2014) or Alzheimer’s disease (Jansen et al., 2019), a specific pathway related to neuronal properties (Kepecs and Fishell, 2014) as in our first HSE example, genetic variants related to brain volume (Jansen et al., 2020), etcetera. The result of such pre-selected GOI is that most of them will likely be related to processes related to the brain, and as such reflect genes likely over-expressed in brain tissue. As a consequence, we argue that in these cases a null condition generated by randomly selecting genes from the total set of ∼20,000 genes (including genes related to all body processes, certainly not only to the brain) is not fair and too liberal, as the null-condition will include a body of background genes not, or less, expressed in brain tissue. Including such genes into the null-condition will lead to a too liberal null-distribution of effects and with that an overestimation of the original effect (de Leeuw et al., 2018).

To avoid this, we advise using a null model that tests whether the observed effect size is larger than the null distribution of effect sizes based on background genes that are particularly expressed in brain tissue. We refer to this null model as “null-brain-gene” (Figure 1c) (Wei et al., 2019), with random genes now selected from the pool of genes that are significantly more expressed in brain tissues in contrast to other body sites. This selection can for example be made according to the GTEx database (Consortium, G. TEx, 2015), containing gene expression data of all sorts of body tissues, including the brain (2957 brain-expressed genes selected by *q* < 0.05, FDR correction, one-sided two-sample *t*-test). In our example, using this stricter null model confirms that HSE genes and connectome metrics are stronger associated than expected for randomly selected brain genes (NOS: *z* = 4.445, *p* < 0.001; SD: *z* = 4.283, *p* < 0.001; FC: *z* = 2.698, *p* = 0.007; Figure 2d)

#### Null-brain-coexpression model

As a further step, we merge the null-coexpressed-gene model and null-brain-gene model, investigating a more integrated null model that tests whether the observed effect size is larger than the null distribution of effect sizes based on brain-expressed genes with similar coexpression level conserved (referred to as “null-coexpressed-brain-gene model”). Using this stringent null model still shows significant associations between HSE gene expression and connectivity strength (NOS: *z* = 4.304, *p* < 0.001; SD: *z* = 4.184, *p* < 0.001; FC: *z* = 2.8934, *p* = 0.004).

### Example 2: Alzheimer’s disease risk gene APOE

Our HSE example includes a case in which the main hypothesized effect survived all different statistical evaluations, from liberal linear regression to stricter (i.e., providing needed spatial and gene specificity) null-coexpressed-gene and null-brain-gene models, showing both spatial and gene specificity. In a second example, we show that this however is not always the case, and that the use of a (too) liberal statistical test not testing for spatial and/or gene specificity potentially may lead to false-positive associations. Here, we zoom in on transcriptomic brain patterns of disease-related pathology, another major topic of combined transcriptomic-neuroimaging studies (Romme et al., 2017; McColgan et al., 2018; Freeze et al., 2019). We examine the Apolipoprotein E (*APOE*) gene, widely indicated as a risk gene for Alzheimer’s disease (AD) (Liu et al., 2013). We hypothesize that the cortical gene expression of *APOE* is related to the cortical alterations revealed in patients with AD or other types of dementia (Grothe et al., 2018), and that correlation analysis between transcriptomic and neuroimaging data may provide evidence for such a relationship.

We again start by examining potential overlap in the transcriptional profile of our gene of interest, here *APOE* (Figure 3a), and a neuroimaging phenotype of interest, here cortical grey matter atrophy patterns of 22 brain diseases, including AD, dementia, and others as reported by meta-analyses of the BrainMap voxel-based morphometry (VBM) studies (see Methods). Linear regression analysis first reveals that the pattern of *APOE* gene expression is significantly associated with results of VBM studies reporting on atrophy of brain regions in *i)* dementia (*β* = 0.653, *p* < 0.001), *ii)* AD (*β* = 0.631, *p* < 0.001), *iii)* semantic dementia (*β* = 0.620, *p* < 0.001) and *iv)* frontotemporal dementia (*β* = 0.505, *p* = 0.001; FDR corrected for multiple testing across 22 diseases; Fig 3b,c). This seems to confirm our hypothesis. Analysis seems to reveal additional associations between *APOE* expression and the atrophy pattern of *v)* attention deficit hyperactivity disorder (ADHD) which has been hypothesized as a risk factor of dementia pathology (Callahan et al., 2017) (*β* = 0.349, *p* = 0.008) and vi) bipolar disorder (*β* = 0.335, *p* = 0.010; Figure 3b).

**Figure 3.**
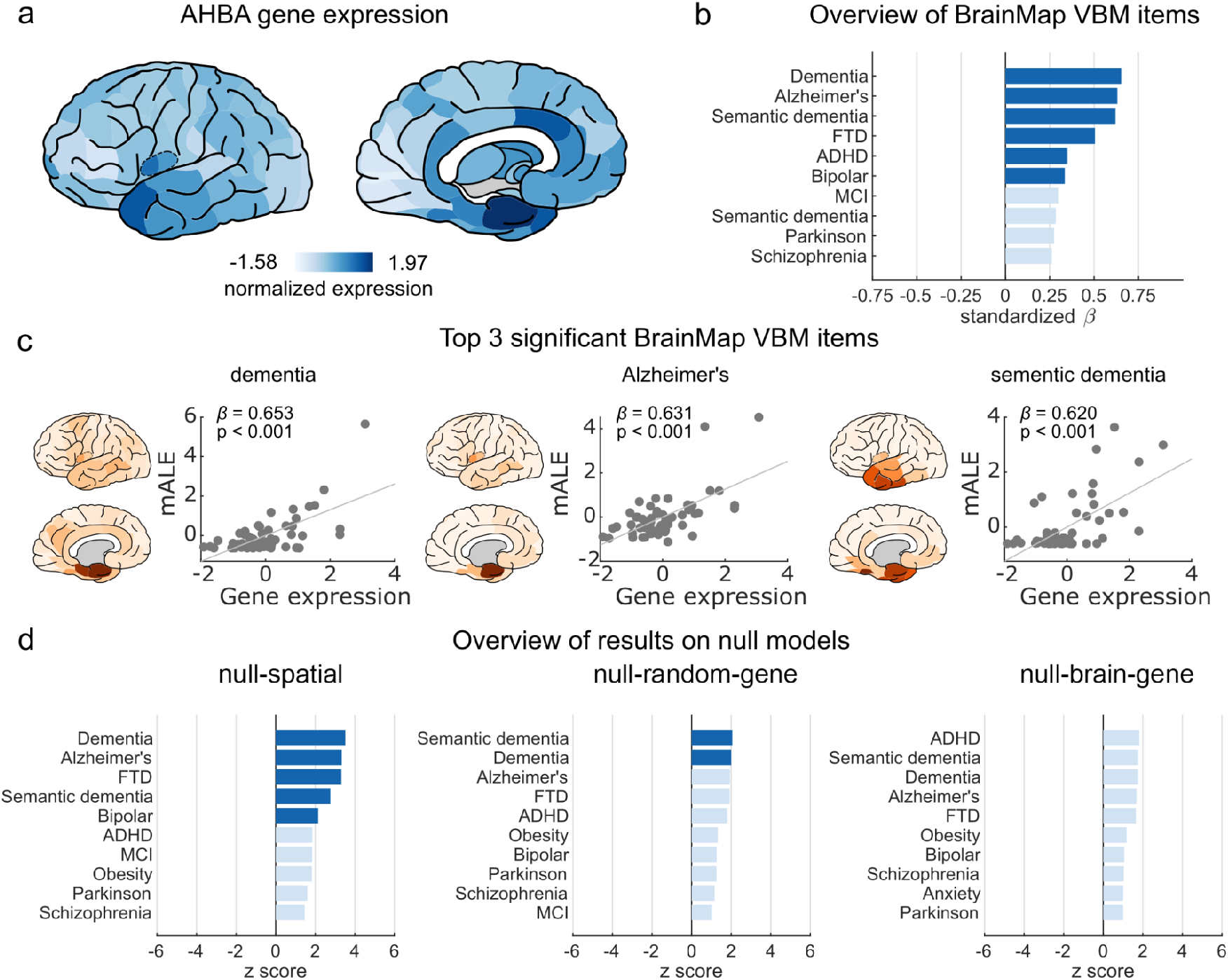
APOE gene expression and atrophy in brain diseases. (a) Brain plots of normalized gene expression levels of *APOE*. (b) Overview of linear regression results, showing the top 10 associations between *APOE* gene expression profile and the BrainMap voxel-based morphometry (VBM)-derived atrophy patterns included in GAMBA. Dark blue indicates significant (*q* < 0.05, FDR corrected across 22 brain diseases included in GAMBA). (c) Top 3 significant correlations between *APOE* gene expression and VBM changes in dementia (*β* = 0.653), Alzheimer’s (*β* = 0.631), semantic dementia (*β* = 0.620). (d) Permutation testing results showing whether the observed effect size (*β* in panel c) is significantly beyond three distinct null distributions of effect sizes for null-spatial, null-random-gene, and null-brain-gene models. Dark blue indicates *p* < 0.05 (two-tailed *z*-test).

As a subsequent step, we use the null-spatial model to examine the *spatial specificity* for the above-mentioned associations between *APOE* transcriptional profile and atrophy patterns in the observed brain diseases. Using this null model reveals significant associations of *APOE*’s expression with brain atrophy in 5 out of the 6 diseases listed above, including dementia (*z* = 3.518, *p* < 0.001), AD (*z* = 3.336, *p* < 0.001), frontotemporal dementia (*z* = 3.331, *p* < 0.001), semantic dementia (*z* = 2.762, *p* = 0.006), and bipolar disorder (z = 2.143, *p* = 0.032; Figure 3d). These findings show encouraging effects that favors a relationship between the transcriptional profile of *APOE* and patterns of cortical atrophy related to dementia.

However, when we evaluate whether the observed associations between *APOE* transcriptional profile and disease-related atrophy patterns exceed effects that one could also observe by chance for *any* other set of genes, these effects diminish. Among the six diseases revealed in linear regression analysis, only the effects for semantic dementia (*z* = 2.057, *p* = 0.040) and dementia (*z* = 2.001, *p* = 0.045) remain significant when using the null-random-gene model; the other effects no longer show significance (AD: *z* = 1.938, *p* = 0.053, frontotemporal dementia: z = 1.928, *p* = 0.054; bipolar disorder: *z* = 1.271, *p* = 0.204). Moreover, using the stricter null-brain-gene model (i.e., zooming in on genes over-expressed in brain tissue) none of the above effects remain significant (all *p* > 0.05). This suggests that the initially found association between the expression pattern of *APOE* and the atrophy patterns of e.g., AD, frontotemporal and semantic dementia, do not exceed effects that can be generally observed when any other (random) gene expressed in brain tissue would have been selected.

### Example 3: Application to autism spectrum disorder risk genes

We further demonstrate the importance of proper null-model selection in avoiding over-interpretation of observed associations in a third example. We examine gene expressions of 25 risk genes of autism spectrum disorder (ASD, 24 of which are present in AHBA) obtained from a recent meta-analysis of genome-wide association studies on a total of 18,381 ASD cases and 27,969 controls (Grove et al., 2019). Again, in a first examination, we compute the correlation between the expression pattern of ASD genes (Figure 4a) and functional alterations in brain diseases derived from BrainMap data (Fox and Lancaster, 2002; Fox et al., 2005; Laird et al., 2005), which reveals an interesting association with Asperger’s syndrome (*β* = 0.284, *p* = 0.032, not corrected) (Figure 4b) -- a major diagnosis of ASD with difficulties in social interaction and nonverbal communication. The subsequent analysis further suggests that this effect for ASD genes is spatially specific as shown by the application of the null-spatial model (*z* = 1.991, *p* = 0.047). However, using null models to examine gene specificity does not show any significant result (null-random-gene: *z* = 1.217, *p* = 0.223; null-coexpressed-gene: *z* = 1.354, *p* = 0.076; null-brain-gene: *z* = 1.246, *p* = 0.213; null-coexpressed-brain-gene: *z* = 1.235, *p* = 0.217) (Figure 4c). This simple example shows that even when an expected association is observed that fits with current theories of brain (dis)organization and (dis)connectivity, these effects may be overestimated, with other sets of genes (in this case, random subsets of genes generally expressed in brain tissue) showing similar expression patterns. We thus again argue that the selection of a proper null model matching one’s research question is of importance in linking brain expression data to neuroimaging phenotypes.

**Figure 4.**
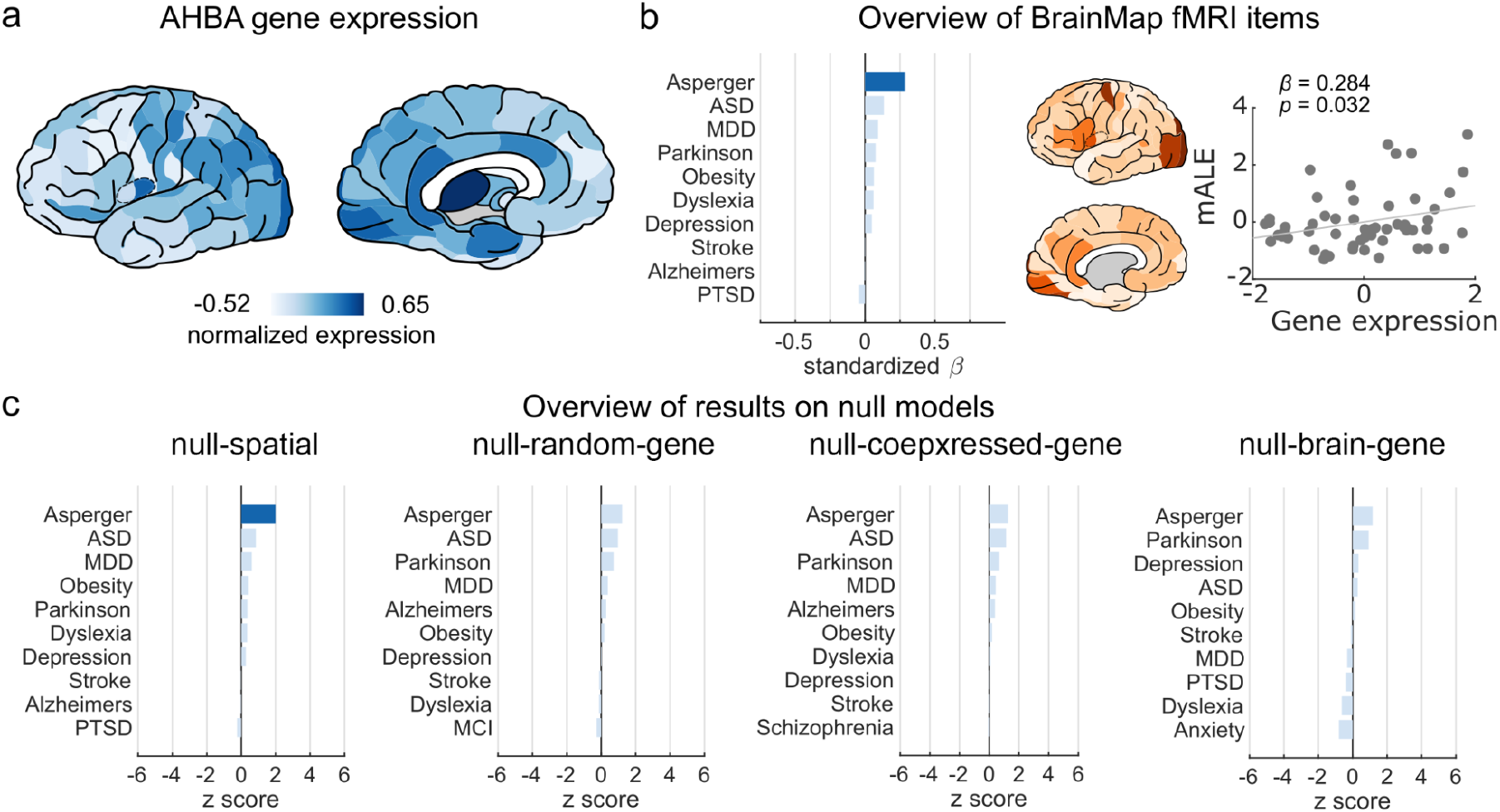
ASD gene expressions and functional alterations in brain diseases. (a) Brain plots of *z*-scores, showing to what extent the expression level of 24 ASD genes is higher than the average gene expression. Yellow circles indicate *p* < 0.05. (b) Overview of linear regression results, showing the top 10 associations between *ASD* gene expression profile and the disease-related BrainMap functional alterations included in GAMBA, with Asperger’s syndrome showing the highest correlation (*β* = 0.284, *p* = 0.032, not corrected). (c) Permutation testing results showing whether the observed effect size (*β* in panel c) is significantly beyond four distinct null distributions of effect sizes for null-spatial, null-random-gene, null-coexpressed-gene, and null-brain-gene models. Dark blue indicates *p* < 0.05 (two-tailed *z*-test).

### Quantitative simulations of null models

#### Brain phenotypes

We provided three simple examples of analyses commonly performed in transcriptomic-neuroimaging studies. We continue by simulating the outcome of the discussed statistical evaluation approaches for a wide range of real brain phenotypes and artificial gradients to get a deeper insight into the comparison of the different null-model strategies and their effect on reporting potential false-positive results. We first build a full outcome-space of all associations between single-gene expression profiles in AHBA (20,949 in total) and 384 imaging-derived brain phenotypes taken from multiple sources such as NeuroSynth (Yarkoni et al., 2011), BrainMap (Fox and Lancaster, 2002; Fox et al., 2005; Laird et al., 2005) and Cognitive Ontology (Yeo et al., 2016) (see methods). Among all these gene × brain phenotype (20,949 × 384) associations, the linear model indicates 960,963 (11.85%) of these associations as significant (*p* < 0.05, uncorrected), with 208,190 (2.57%) reaching FDR and 56,259 (0.69%) reaching Bonferroni correction with *α* level < 0.05 (corrected for 384 tests of all brain phenotypes per gene). We then implemented the null models to examine the spatial/gene specificity for the potential significant associations. Only FDR-corrected results are reported in the following paragraphs for the sake of simplicity. Uncorrected and Bonferroni-corrected results are tabulated in Table 2.

**Table 1.**
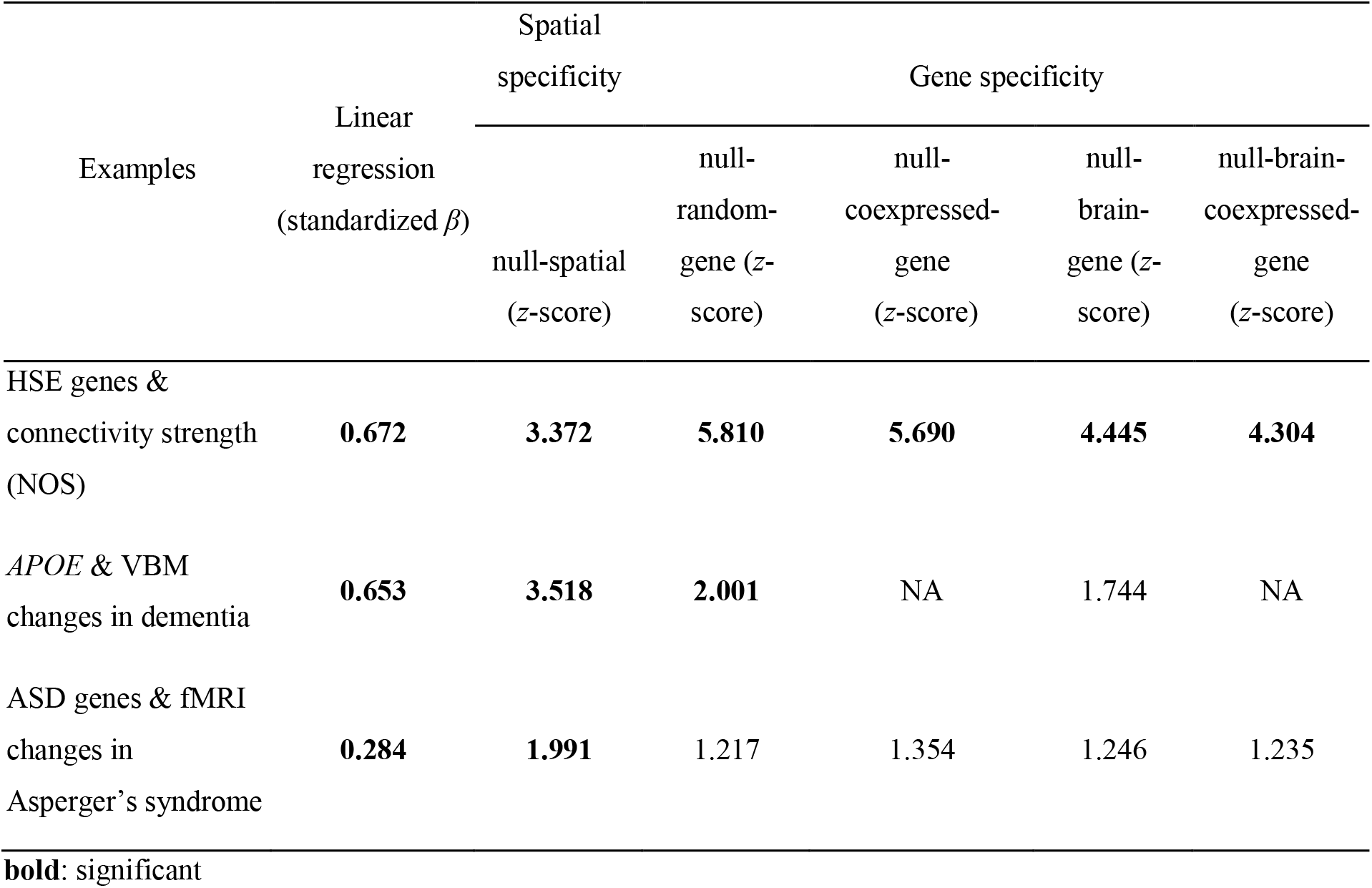
Summary of examples statistically testing associations between spatial patterns of gene expression and imaging-derived brain phenotypic patterns

**Table 2.** Summary of all potential associations between spatial patterns of single-gene expression and 384 imaging-derived brain phenotypic patterns

| Methods | Number of significant associations ($\alpha < 0.05$) | | |
| --- | --- | --- | --- |
|  | Uncorrected | FDR-corrected | Bonferroni-corrected |
| LR | 960,963 | 208,190 | 56,259 |
| LR & N-spin | 84,149 | 26,273 | 8,142 |
| LR & N-spin & N-rand | 26,786 | 15,297 | 7,610 |
| LR & N-spin & N-brain | 19,551 | 9,802 | 6,294 |

LR: Linear Regression; N-spin: null-spatial model; N-rand: null-random-gene model; N-brain: null-brain-gene model

**Table 3.** Summary of all potential associations between spatial patterns of single-gene expression and 7 simulated brain phenotypes that represent global spatial gradients

| Methods | Number of significant associations ($\alpha < 0.05$) | | |
| --- | --- | --- | --- |
|  | uncorrected | FDR-corrected | Bonferroni-corrected |
| LR | 33,335 | 23,723 | 18,336 |
| LR & N-spin | 3,880 | 3,305 | 3,175 |
| LR & N-spin & N-rand | 1,003 | 974 | 967 |
| LR & N-spin & N-brain | 647 | 626 | 620 |

LR: Linear Regression; N-spin: null-spatial model; N-rand: null-random-gene model; N-brain: null-brain-gene model.

Using the null-spatial model to assess spatial specificity of these associations shows that only 26,273 out of the 208,190 associations reported in linear regression remain significant (*p* < 0.05; Supplementary Figure S1). This suggests that a large proportion of transcriptomic-neuroimaging associations as revealed by linear regression (87%) are overestimated due to dependencies of expression levels among neighboring brain regions and thus likely involve false-positive findings. This is in line with what was recently reported in literature (Burt et al., 2020; Fulcher et al., 2020; Markello and Misic, 2021).

When we then further implement the null-random-gene and null-brain-gene models (null-coexpressed-gene is not applicable in the situation of examining the spatial pattern of a single gene) to examine gene specificity of the reported transcriptomic-neuroimaging associations of our gene of interest and all neuroimaging patterns, we can find that only 15,297 out of these 26,273 reported associations surviving the null-spatial model remain further significant using the null-random-gene (*p* < 0.05, null-random-gene and null-spatial combined). This suggests that even among spatially specific associations, there is still a considerable proportion of associations (42%) that can be commonly found for a wide range of (random) genes and therefore are hard to call an effect that is specific to our gene of interest. These effects thus likely reflect effects that are not specific to the gene of interest and reflect patterns that are found for many other genes as well. Further examining effects under the more stringent null-brain-gene model shows that an even smaller proportion of only 9,802 associations out of the 26,273 associations surviving the null-spatial model remain significant (*p* < 0.05, null-brain-gene and null-spatial combined; Supplementary Figure S1).

We note that similar types of results are found when we apply the null models the other way around, namely, first applying the null-brain-gene model and then the null-spatial model. Doing so, we can identify 67,181 significant transcriptomic-neuroimaging associations (*p* < 0.05, null-brain-gene), of which only 15% (9,802) are spatially specific (*p* < 0.05, null-spatial and null-brain-gene combined; Supplementary Figure S1), indicating that gene and spatial specificity are not mutually inclusive.

Similar findings can also be observed for gene sets of more than one gene by examining transcriptomic-neuroimaging associations for gene sets curated in the Gene Ontology (GO) database (http://geneontology.org) (results for GO terms are presented in the Supplementary Results).

#### Simulated phenotypes

In the previous paragraphs we demonstrate the need of examining spatial and gene specificity of observed transcriptomic-neuroimaging associations. We argue that this is particularly important when testing phenotypes with spatial patterns similar to global posterior-anterior or inferior-superior gradients (McColgan et al., 2018; Vogel et al., 2020). To show this point, we simulated spatial maps of seven phenotypes that follow global spatial gradients across the brain, from i) posterior to anterior, ii) from inferior to superior, and iii) from medial to lateral, together with the (iv-vii) four combinations of these gradients (Supplementary Figure S2). We then examined all associations between single-gene expression profiles in AHBA and the simulated phenotypes. Among all these possible associations (20,949 × 7), we can find 23,723 (16%) significant associations between single-gene transcriptional profiles and simulated phenotypic profiles (linear regression: *q* < 0.05, FDR corrected; corrected for 7 tests of all simulated phenotypes per gene). These correlations suggest that a large set of genes have expression profiles that follow general and very global spatial gradients of the brain, and show a non-specific spatial pattern that is shared with a wide range of other genes (i.e., lack any form of gene specificity). Application of the null-spatial model to these associations shows that only 3,305 out of the observed 23,723 associations (14%) remain significant, suggesting that most of the common effects found significant using linear regression are inflated due to the strong spatial dependency of neighboring brain regions and can be filtered out by the null-spatial model. Among those 3,305 spatial-specific associations however, only 974 (29%) further remain significant when we additionally apply the null-random-gene model and examine whether effects are also gene specific. An even smaller number of significant associations remain (only 626; 19%) when we use a stricter null-brain-gene model (Supplementary Figure S1).

These simulations show that only 3% (626 out of 23,723) of the associations between single-gene transcriptional profiles and 8 simple spatial gradients revealed in ordinary linear regression can be labeled as both spatially specific and gene specific; 97% of the initially found FDR-corrected effects do not survive a statistical evaluation that examines spatial and gene specificity.

### Toolbox

We made a simple web-based application and a MATLAB toolbox to facilitate quick examinations of transcriptomic-neuroimaging associations and to then test observed correlations with different null models. Within GAMBA (short for Gene Annotation using Macroscale Brain-imaging Association) expression profiles of input GOIs (i.e., a single gene or a set of genes) can be associated to imaging-derived brain traits from nine categories (Supplementary Figure S3) and tested using the different null models as discussed in this paper (including null-spatial, null-random-gene, null-coexpressed-gene, null-brain-gene). Imaging-derived phenotypes included in the tool cover the spatial patterns of *i*) resting-state functional networks (Yeo et al., 2011), *ii*) brain cognitive components (Yeo et al., 2016), *iii*) regional metrics of the brain structural and functional connectome, *iv*) measurements of the cortical oxygen and glucose metabolism (Vaishnavi et al., 2010), *v*) human surface area expansion compared to the chimpanzee (Wei et al., 2019), *vi*) brain volume alterations across twenty-two disorders (Fox and Lancaster, 2002; Fox et al., 2005; Laird et al., 2005), *vii*) brain functional changes in sixteen disorders (Fox and Lancaster, 2002; Fox et al., 2005; Laird et al., 2005), *viii*) cortical patterns of brain (dis)connectivity across nine psychiatric and neurological disorders (de Lange et al., 2019) and *ix*) brain functional correlates of 292 terms in relation to cognitive states and brain disorders, as described in the NeuroSynth database (Yarkoni et al., 2011). Details on the included datasets and statistical analyses are described in the Supplementary Methods. The web-tool is available online at http://dutchconnectomelab.nl/GAMBA. A relevant MATLAB toolbox used for implementing the mentioned null models is also available at https://github.com/dutchconnectomelab/GAMBA-MATLAB.

## Discussion

We evaluate and discuss the importance of selecting proper null conditions when performing a transcriptomic-neuroimaging study. We examined the usability of commonly applied statistical methods, such as simple single linear regression to assess the statistical validity of observed transcriptomic-neuroimaging relationships and once again confirm that the use of proper null-models that provide spatial specificity (using null-spatial model) are highly needed. We further show, however, that preserving spatial effects and controlling for spatial autocorrelation is not enough to provide information on whether the effect of our gene(s) of interest are unique and stand-out from effects that can be widely observed across many other random genes in the dataset. We recommend the use of appropriate null models when examining overlapped spatial patterns of transcriptomic profiles and imaging-derived phenotypes, providing information on both spatial and gene specificity based on the research question asked.

The three examples we present do of course not cover all types of examinations one can perform by combining transcriptomic and neuroimaging data. They do however illustrate some of the caveats and limitations of cross-linking cortical patterns of transcription and neuroimaging features and recommend the use of additional statistical analyses. Our first example illustrates that transcriptomic-neuroimaging associations can include effects that are both spatial- and gene-specific. We show that the cortical transcriptional profile of HSE genes runs parallel to spatial patterns of macroscale brain traits and that this association goes beyond both general spatial and random-gene-set effects that could be expected based on chance level alone. They thus highlight an interesting relationship between expression patterns of genes related to patterns of macroscale connectivity (Zeng et al., 2012; Krienen et al., 2016; Romero-Garcia et al., 2018b).

A different conclusion should however be made when examining *APOE* and the set of ASD genes, with the two examples underscoring the importance of proper statistical testing. The expression pattern of *APOE* across cortical areas seems to show, at first glance, a sensible and significant correlation with reported patterns of structural alterations in AD and dementia. Findings in literature are in support of such associations, arguing in favor of a “true positive” effect. For example, AD patients are known to show elevated *APOE* expression in the medial temporal regions that are in turn known to be involved in the pathology of the disease (Akram et al., 2012; Linnertz et al., 2014; Ranlund et al., 2018). However, further evaluation using null models that test gene specificity reveals that the observed association is not particularly specific to *APOE*, and can be observed for nearly 150 other brain-expressed genes in AHBA (corresponding to *z* = 1.65 for AD, null-brain-gene model). This indicates that while potentially an interesting effect, it is quite hard to say that this effect is specific to *APOE*. Investigating a gene set involved in AD-related biological pathways instead of a single risk gene may produce more informative results (Freer et al., 2016).

Similarly, analysis of ASD risk genes initially reveals an interesting association between expression patterns of ASD genes and patterns of brain functional alterations seen in Asperger’s syndrome, as shown by both linear regression and the null-spatial model. One may be easily tempted to take this effect as a meaningful interaction because Asperger’s syndrome is a highly heritable subtype of ASD (Grove et al., 2019) and imaging-genetic associations in ASD have been proposed (Ranlund et al., 2018; Xie et al., 2020). Yet, our analyses using null-random-gene and null-brain-gene models show that we cannot claim specificity of this effect to ASD genes, as such an overlapping pattern can similarly be found for (many) other random genes not related to ASD. This example indicates that a false-positive conclusion can easily be made and that using null models to test gene specificity is important.

Simulations correlating single-gene expression profiles to brain phenotypes show that actually only 13.6% of all observed linear associations that survive FDR satisfy spatial specificity. Such a small proportion is likely attributable to a high auto-correlation between adjacent brain regions in transcriptomic and phenotypic data, leading to a degree of freedom much smaller than the one applied, such that the effects are inflated. This again stresses the importance of using null models [e.g., the spin-based null-spatial model here (Alexander-Bloch et al., 2018) or other equivalents (Arnatkevic̆iūtė et al., 2019; Burt et al., 2020; Fulcher et al., 2020; Markello and Misic, 2021)] to reduce false-positive rate introduced by the spatial auto-correlation effects.

Extending the important null-spatial model, our simulation analysis correlating single-gene expression profile to brain phenotypes shows that implementing the null-spatial model only is, however, not enough, and that only 37.3% of the transcriptomic-neuroimaging associations survived in the null-spatial model do not show *gene specificity*. The ratio is even lower (18.9%) in our simulations that take brain geographic gradients as the phenotypic of interest. This is because the transcriptional profiles of a considerable number of genes (16% of all genes) bear high similarity to the examined geographic gradients in the brain. These simulations suggest that examining spatial specificity cannot be regarded as a substitute of testing gene specificity, nor vice versa, pointing to the necessity of testing both the spatial and gene specificity in transcriptomic-neuroimaging studies.

The discussed null models are presented in a simple web-tool GAMBA and a MATLAB toolbox, which can be used to probe the association between transcriptomics and common brain structure/function phenotypes derived from a wide range of neuroimaging data. GAMBA complements other tools that link genetics and brain functions, tools such as NeuroSynth (Yarkoni et al., 2011), Brain Annotation Toolbox (Liu et al., 2019), and the ENIGMA toolbox (Larivière et al., 2021) that provide similar platforms to visualize and examine brain maps of gene expression patterns and brain maps, but do not directly provide means to test these effects against multiple null models.

Several methodological points have to be considered. First, our examples are based on the rich gene expression data from the AHBA (Hawrylycz et al., 2012). Further extensions to other datasets of brain gene expressions, such as PsychENCODE (Wang et al., 2018) and BrainSpan (Johnson et al., 2009) can easily be made. Second, we need to consider that AHBA gene expression data is extracted from the post-mortem brains of healthy individuals, i.e., ones with no psychiatric and neurologic conditions. Therefore, results regarding spatial brain patterns of disorders should be interpreted in the context of gene expression of risk genes in brain regions to be a potential marker for a higher susceptibility of these regions in relevant disorders (Romme et al., 2017; McColgan et al., 2018).

## Conclusion

In summary, we highlight the need of using null models to investigate both spatial specificity and gene specificity when examining associations between the spatial patterns of gene transcriptomic profiles and imaging-derived brain traits.

## Methods

### AHBA gene expression data

Microarray gene expression data were obtained from the extensive Allen Human Brain Atlas database (http://human.brain-map.org), including highly detailed data from six postmortem brains of donors without any neuropathological or neuropsychiatric conditions. Microarray analyses are described in detail in http://help.brain-map.org/display/humanbrain/Documentation. Brain tissue samples of the left hemisphere were obtained from four donors (466 ± 72.6 samples from H0351.1009, H0351.1012, H0351.1015, and H0351.1016), and 946 and 893 samples covering both hemispheres from the remaining two donors (H0351.2001 and H0351.2002). We included tissue samples of cortical and subcortical regions of the left hemisphere and used the expression of 58,692 probes for each brain donor (Romme et al., 2017; Wei et al., 2019).

We performed probe-to-gene re-annotation using the BioMart data-mining tool (https://www.ensembl.org/biomart/) (Arnatkevic̆iūtė et al., 2019). Outdated gene symbols were updated and alias gene symbols were replaced by symbols obtained from the HUGO Gene Nomenclature Committee (HGNC) database (http://biomart.genenames.org/), resulting in the inclusion of 20,949 genes. Expression levels that were not well-above background were set to *NaN*. Per donor, and per tissue sample, expression levels of probes annotated to the same gene symbol were averaged, followed by log2-transformation with pseudocount 1. Tissue samples were spatially mapped to FreeSurfer cortical and subcortical regions to obtain region-wise gene expression profiles (French and Paus, 2015). A cortical parcellation of 114 regions (57 per hemisphere) and subcortical segmentation of 7 regions based on the Desikan-Killiany atlas (DK-114) were obtained for the Montreal Neurological Institute (MNI) 152 template using FreeSurfer (Desikan et al., 2006; Fonov et al., 2011; Cammoun et al., 2012). Brain tissue samples were annotated to cortical regions in the DK-114 atlas based on MNI coordinates, computing the nearest gray matter voxel within the MNI ICBM152 template in the FreeSurfer space. Tissue samples with a distance less than 2mm to the nearest gray matter voxel were included. Gene expression profiles of tissue samples belonging to the same cortical region were averaged, resulting in a 6 × 64 × 20,949 data matrix (i.e., donors × brain regions × genes). Within each donor, gene expression per gene was normalized to *z-*scores across all cortical and subcortical regions. Normalized gene expression profiles were averaged across the six donors obtaining a group-level gene expression matrix of size 64 × 20,949. Considering that most neuroimaging data as described in the following sections only include cortical regions, only the cortical expression pattern of each gene (i.e., 57 × 20,949 gene expression matrix) was correlated to the patterns of various neuroimaging findings.

### Neuroimaging phenotypic data

Connectome metrics used in *example 1* were obtained from a human structural connectome map reconstructed using T1-weighted and diffusion-weighted MRI (dMRI) data of 487 subjects (age [mean ± standard deviation]: 29.8 ± 3.4 years old) from the Human Connectome Project (Van Essen et al., 2013). Disease maps used *in example 2* and *example 3* were computed based on coordinate-based results obtained from the extensive BrainMap database (http://www.brainmap.org/) that contains published functional and structural neuroimaging experiments of psychiatric and neurological disorders (see Supplementary Methods for a detailed description of the used procedures).

### Statistics and null models

#### Linear regression

Linear regression is used to test for an association between the cortical expression profile of a gene (or the average expression pattern across a set of genes) and the pattern of an imaging-derived brain phenotype:

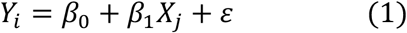

where *Y*_i_ indicates the standardized gene expression profile of gene *i* or the standardized, averaged profile of a gene set *i*, and *X*_j_ the standardized cortical profile of neuroimaging phenotype *j*. Standardization is performed by dividing each value *X* or *Y* by the standard deviation. The standardized regression coefficient *β*_1_ and the corresponding correlation coefficient and *p*-value are obtained.

#### Null-spatial model

An important variant on the linear regression model was recently introduced (Alexander-Bloch et al., 2018), now testing whether the observed association is specific to spatial-anatomical relationships between brain regions. To this end, in (1) the observed *β*_1_ is compared to *β*_1_s generated by 1,000 permutations, in which the gene expression data matrix is rebuilt using randomized brain parcellations by spinning the reconstructed sphere of the real brain parcellation with random angles (0-360°) conserving the spatial relationship of neighboring regions.

#### Null-random-gene model

The null-random-gene model tests whether the observed association is specific to the given gene or set of genes of interest (GOI), i.e., comparing against effects that can be observed when any other set of genes would have been selected. To this end, it is tested whether the observed *β*_1_ (i.e., the effect size) is different from null distributions of *β*_1_ observed for randomly selected genes with the null-random-gene distribution estimated from 10,000 permutations and the mean (*µ*) and standard deviation (*σ*) of *effect sizes* in the null distribution obtained.

#### Null-coexpressed-gene model

The null-coexpressed-gene model includes a stricter null model where random genes with similar co-expression levels as the given GOI are selected to generate null distributions of *β*_1_ (null-coexpressed-gene model). Random genes are selected according to the following steps: first, a set of random genes are initially selected. Then the mean co-expression level of the random gene set is compared to the mean co-expression level of the given GOI. If the co-expression difference is larger than the maximum difference allowed, the gene with the highest/lowest co-expression level is excluded and a new random gene is included in the set. This step repeats until the co-expression difference is smaller than the maximum difference allowed. A total number of 1,000 sets of random genes were obtained due to the limitation of computational capacity.

#### Null-brain-gene model

The null-brain-gene model is generated using random genes selected from a pool of 2,957 genes over-expressed in brain tissue, with brain-expressed genes identified by performing one-tail two-sample t-tests on gene expression levels between brain tissues and other body sites (*q* < 0.05, FDR corrected), using gene expression data from the GTEx portal (https://www.gtexportal.org). Permutation is performed 10,000 times.

For all null models, the mean (*µ*) and standard deviation (*σ*) of effect sizes (*β*_1_) in the null distributions are estimated. A two-tailed *z*-test is performed to examine whether the observed *β*_1_ (i.e., the effect size) is larger than the mean effect size derived from null models:

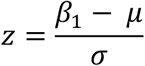

where *µ, σ* indicate the mean and standard deviation of *β*_1_ over random permutations. A two-tailed *p*-value is computed as follows:

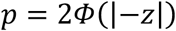

where Φ is the standard normal cumulative distribution function.

## Data availability

Gene expression data are available in the Allen Brain Atlas (http://human.brain-map.org). The GTEx data are available in the GTEx Portal (https://www.gtexportal.org). MRI data (used for the connectome reconstruction) that support the findings of this study are available from the Human Connectome Project (Q3 release, https://www.humanconnectome.org). BrainMap data are available at http://www.brainmap.org/.

## Author contributions

**Y**.**W**.: methodology, software, investigation, writing - original draft, writing - review & editing; **S**.**C**.**d**.**L**.: methodology, software, writing - review & editing; **R**.**P**.: validation, writing - review & editing; **L**.**H**.**S**.: visualization, writing - review & editing; **DJ**.**A**.: visualization, writing - review & editing; **K**.**W**.: software, writing - review & editing; **D**.**P**.: Supervision, writing - review & editing; **M**.**P**.**v**.**d**.**H**.: conceptualization, writing - review & editing, supervision;

## Acknowledgments

The work of M.P.v.d.H. was supported by an ALW open (ALWOP.179) and VIDI (452-16-015) grant from the Netherlands Organization for Scientific Research (NWO) and a Fellowship of MQ. D.P. was supported by The Netherlands Organization for Scientific Research (NWO VICI 453-14-005). S.C.d.L. was supported by the Amsterdam Neuroscience alliance grant. Human neuroimaging data was kindly provided by the Human Connectome Project, WU-Minn Consortium (Principal Investigators: David Van Essen and Kamil Ugurbil; 1U54MH091657) funded by the 16 NIH Institutes and Centers that support the NIH Blueprint for Neuroscience Research; and by the McDonnell Center for Systems Neuroscience at Washington University.

