## Supplementary Materials for "Statistical testing in gene transcriptomic-neuroimaging associations: an evaluation of methods that assess spatial and gene specificity"

### Supplementary Methods

*Neuroimaging phenotypes in GAMBA*

*Connectome metrics.* Connectome metrics were obtained from a human structural connectome map reconstructed using T1-weighted and diffusion-weighted MRI (dMRI) data of 487 subjects (age [mean ± standard deviation]: 29.8 ± 3.4 years old) from the Human Connectome Project (Van Essen et al., 2013). White matter tracts were reconstructed from the dMRI data using deterministic tractography (Wei et al., 2019b). The 18 sets of b = 0 volumes were averaged, and the 270 diffusion images were realigned and corrected for motion and gradient-induced distortions (Andersson and Skare, 2002). The diffusion profile within each voxel was reconstructed using generalized q-sampling imaging (GQI) (Yeh et al., 2010) and deterministic tractography was used to reconstruct white matter tracts, performing fiber assignment using the Fiber Assignment by Continuous Tracking (FACT) algorithm (Mori et al., 1999). Eight streamline seeds were started per voxel and tracking was stopped if the streamline reached a voxel of low preferred diffusion direction (fractional anisotropy [FA] < 0.1), exited the gray matter/white matter mask, or made a sharp turn (> 45°). Next, a structural connectome was generated from the set of reconstructed tractography streamlines and the cortical parcellation of each subject. Desikan-Killiany (DK) 114 atlas was used to parcellate the cortex into 114 regions (57 per hemisphere). The number of streamlines (NOS), streamline density (SD), mean fractional anisotropy (FA) were extracted and used as the weight of structural connectivity between cortical regions. A group-averaged binary network was formed by averaging values across subjects for connections observed in more than 60% of the subjects (de Reus and van den Heuvel, 2013). Nodal degree was calculated as the number of connections linked to each region. Nodal strength was computed for each cortical region by summing up the weights of all connections linked to that region.

A functional connectome map was constructed using resting-state fMRI data. Data were realigned and co-registered with the T1-weighted image to overlap with the cortical parcellation maps. Blood oxygenation level-dependent (BOLD) time series were corrected for linear trends, as well as global nuisance covariance, including six head motion parameters and mean signals of white matter and the ventricles. We performed band-pass filtering (0.01-0.1 Hz) together with motion scrubbing to minimize the influence of head motion (Power et al., 2012). Regional time series were computed from the preprocessed fMRI data by averaging the time series of all voxels within a cortical region. Interregional functional connectivity was calculated as the Pearson's correlation coefficient between the time series of cortical regions. Negative correlations were excluded, and correlation coefficients were transformed to $z$-scores using Fisher transformation. A group-level functional network was constructed by averaging $z$-scores across all subjects. Nodal strength of functional connectivity was subsequently computed by summing up the weights of functional connectivities linked to that region.

*Anatomical and functional involvement in brain diseases.* The extensive BrainMap database (http://www.brainmap.org/) containing published functional and structural neuroimaging experiments with coordinate-based results was used to extract brain mappings of a wide range of psychiatric and neurological disorders. MNI coordinates related to 22 brain disorders from the voxel-based morphometry (VBM) database (Number of experiments = 46 ± 48, ranging from 4 to 206) and of 16 brain disorders from the functional database (Number of experiments = 69 ± 69, ranging from 13 to 267) were obtained using Sleuth (Fox et al., 2005; Fox and Lancaster, 2002; Laird et al., 2005; Vanasse et al., 2018). Meta-analysis was performed for each disorder using activation likelihood estimation (ALE) implemented in GingerALE (Eickhoff et al., 2012; Turkeltaub et al., 2012). ALE maps were registered to the

MNI152 template in the FreeSurfer space using FLIRT (Jenkinson et al., 2002; Jenkinson and Smith, 2001). Furthermore, ALE scores of gray matter voxels within each DK region were averaged, resulting in two matrices of size 114 × 22 and 114 × 16, indicating the respective anatomical and functional cortical involvement of each disorder.

*Functional networks.* Cortical regions in the DK-114 atlas were assigned to seven well-defined functional networks using the Yeo 7-network atlas (Yeo et al., 2011), including the visual, somatomotor, dorsal-attention, ventral-attention, limbic, frontal-parietal, and default-mode network. Surface-based annotation of the functional networks on the averaged subject in FreeSurfer (i.e., *fsaverage*) was translated to a 3D brain image in volumetric space, in which each gray matter voxel was annotated to the closest functional network. Per DK region the ratio of voxels that belonged to each of the seven functional networks was computed, resulting in a 114 × 7 matrix mapping the involvement of each DK region in each of the functional networks.

*Brain cognitive components.* Brain regions were mapped to the functional ontology of twelve distinct brain cognitive components from https://surfer.nmr.mgh.harvard.edu/fswiki/BrainmapOntology_Yeo2015 and translated to 3D brain images in volumetric space. These components relate to various cognitive and behavioral processes based on functional neuroimaging data from 10,449 experiments (Yeo et al., 2016). A mapping matrix of size 114 × 12 was computed indicating the level of involvement of each of the 114 cortical areas in each of the twelve cognitive components.

*Metrics of cortical metabolism.* Data of five cortical metabolic measurements obtained from positron emission tomography (PET) imaging (tabulated and described in detail in (Vaishnavi et al., 2010)) was used, including the vertebral metabolic rate for oxygen ($CMRO_2$), cerebral metabolic rate for glucose (CMRGlu), glycolytic index (GI), oxygen-glucose index (OGI), and cerebral blood flow (CBF). Data of metabolic measurements were given based on the Brodmann atlas (BA) and were transformed to the DK-114 atlas using the following steps. MNI coordinates of all gray matter voxels of the *fsaverage* subject in FreeSurfer were extracted and MNI coordinates were mapped to the Talairach atlas to obtain the annotation of each BA area, using the BioImage Suite (https://bioimagesuiteweb.github.io/webapp/mni2tal.html) (Lacadie et al., 2008). With the resulting annotations, we assigned the value of metabolic measurements to each voxel in the gray matter of the *fsaverage* subject. For each region of the DK atlas mean values were calculated per metabolic measurement, among all gray matter voxels within that region, yielding a 114 × 5 data matrix of metabolic measurements.

*Metric of evolutionary cortical expansion.* A brain map of evolutionary cortical expansion was obtained from (Wei et al., 2019a), where T1-weighted MRI data from 29 chimpanzees and 50 humans were utilized to compute the surface area (*S*) expansion of each homologous cortical region in the DK-114 atlas. The expansion was computed as $(S_{human} - S_{chimpanzee}) / S_{chimpanzee}$.

*NeuroSynth functional neuroimaging metrics.* The comprehensive NeuroSynth database was used to obtain brain mappings of 749 terms, with voxel-wise *z*-scores representing the extent to which a voxel is associated with the corresponding term (FDR corrected, $q < 0.01$). Brain terms and matching images were generated using text-mining, meta-analysis and machine-learning techniques on fMRI studies mapping the involvement of brain regions in a large variety of cognitive tasks and states (http://www.neurosynth.org) (Yarkoni et al., 2011).

Images of 457 terms with abstract meaning (e.g., "association"), or with names of brain anatomy (e.g., "putamen") were excluded. Resulting 292 images (listed in Supplementary Table 3) were registered to the MNI152 template in FreeSurfer space using FLIRT (Jenkinson et al., 2002; Jenkinson and Smith, 2001). The averaged *z*-score within each cortical region of the DK-114 atlas was computed, resulting in a 114 × 292 matrix to indicate the cortical involvement in each term.

### Supplementary Results

*GO gene set and imaging-derived brain phenotypes*

We examined the number of potential associations in terms of different statistical evaluation methods for 7,350 gene sets that are related to biological processes curated in the Gene Ontology (GO) database (http://geneontology.org). Among all 7,350 × 384 examinations, we noted 255,682 potential associations (30,001 reaching FDR correction, and 9,027 reaching Bonferroni correction) using simple linear regression. A major proportion of these associations however were flagged as potential false-positive effects due to the effects do not exceed what one can expect in the null-spatial model (93.35% of the uncorrected, 90.65% of the FDR-corrected, and 91.97% of the Bonferroni-corrected results), remaining 17,002 (uncorrected), 2,806 (FDR corrected), and 725 (Bonferroni corrected) significant associations. Furthermore, examining gene specificity by means of the null-random-gene model showed that 1,545 out of the 16,920, 194 out of the 2,806, and 30 out of the 725 observed associations may be no specific to the input gene set. Using the null-coexpressed-gene model revealed more associations [1624 (uncorrected), 205 (FDR-corrected), 34 (Bonferroni-corrected), respectively] that could not show gene specificity. Using the stricter null-brain-gene model revealed more associations [1870 (uncorrected), 237 (FDR-corrected), 42 (Bonferroni-corrected), respectively] that could not survive the null-brain-gene model, suggestive of effects that were not specific to the gene set of interest. These results suggest that it is necessary to implement both the null-spatial model and null-brain-gene model to avoid potential false-positive findings of transcriptomic-neuroimaging associations.

*Over-expression of the GOI*

An important application of brain transcriptomic data is to study where in the brain the gene(s) of interest (GOI) is (over)expressed, an analysis often performed by evaluating the expression level of the GOI in comparison to the mean expression of all genes in that area (Jansen et al., 2019).

In our HSE example, up-regulated expression of HSE genes is found within the inferior/superior parietal cortex, supramarginal cortex, precuneus, caudal frontal cortex, precentral gyrus, and paracentral cortex [two-tailed $z$-test; $q < 0.05$, False Discovery Rate (FDR) corrected for multiple comparisons across 64 tests (i.e., within 57 cortical + 6 subcortical regions); Supplementary Figure S4]. Gene expression levels of HSE genes in the above regions are significantly higher than the rest of the brain (two-tailed two-sample $t$-test, $t = 6.617$, $p < 0.001$), suggesting over-expression of HSE genes in these brain regions specifically.

In the examples of *APOE* and ASD genes, we however do not find any significant over-expression across brain regions. Examining the spatial location of the highest expression level of *APOE* across the brain shows the entorhinal cortex to be among the highest expressed regions ($z = 2.215$, $p = 0.013$; not significant after FDR correction for multiple testing across 64 brain regions; Supplementary Figure S4). ASD genes show higher-than-average expression in the lingual gyrus ($z = 2.591$, $p = 0.010$) and lateral occipital lobe ($z = 2.573$, $p = 0.010$; not significant after FDR correction for multiple testing across 64 brain regions; Supplementary Figure S4).

### Supplementary Figures

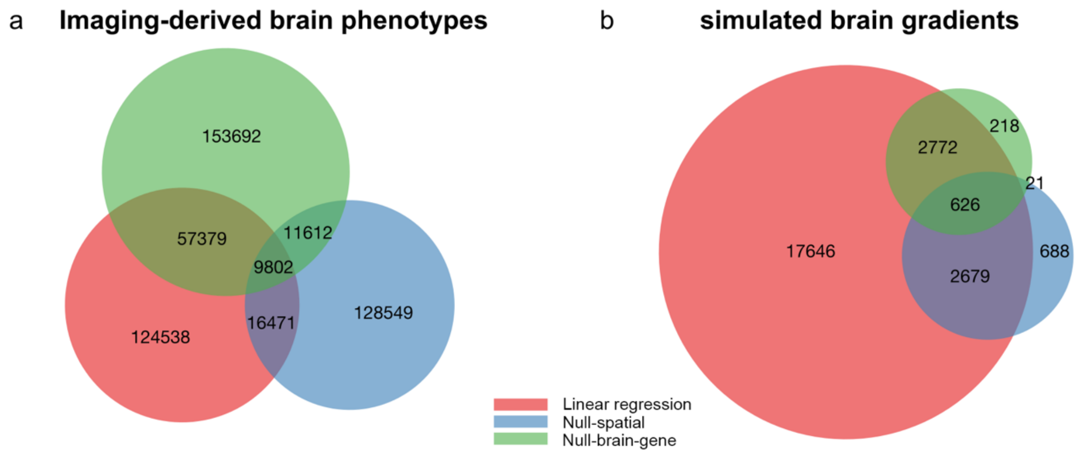


Figure S1. Venn diagram of the number of significant associations between single-gene transcriptional profiles and (a) 384 imaging-derived brain phenotypic patterns included in GAMBA and (b) 7 global spatial gradients in the brain.

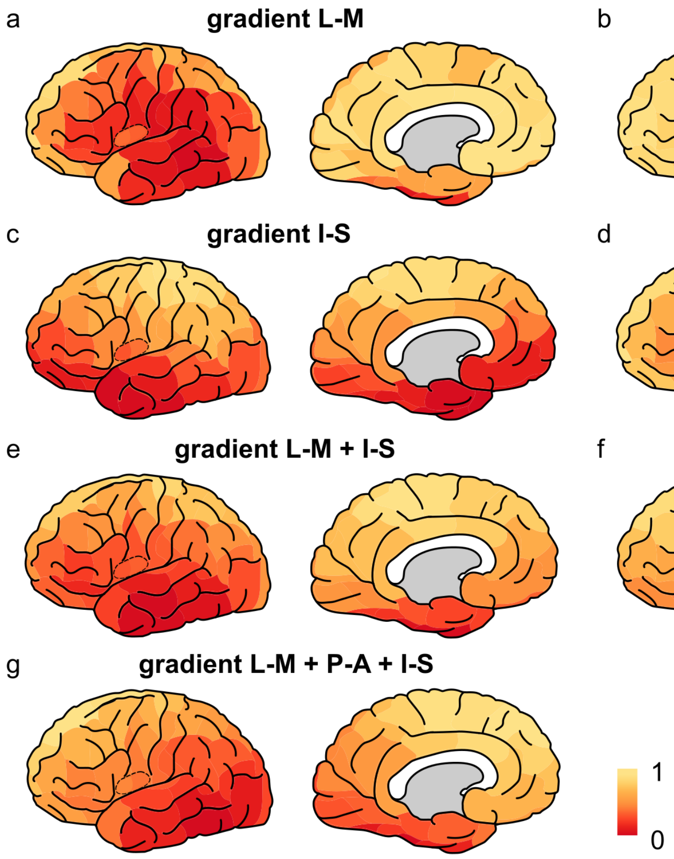


Figure S2. Brain maps of the seven global spatial gradients in the brain. L-M: lateral-medial; P-A: posterior-anterior; I-S: inferior-superior.

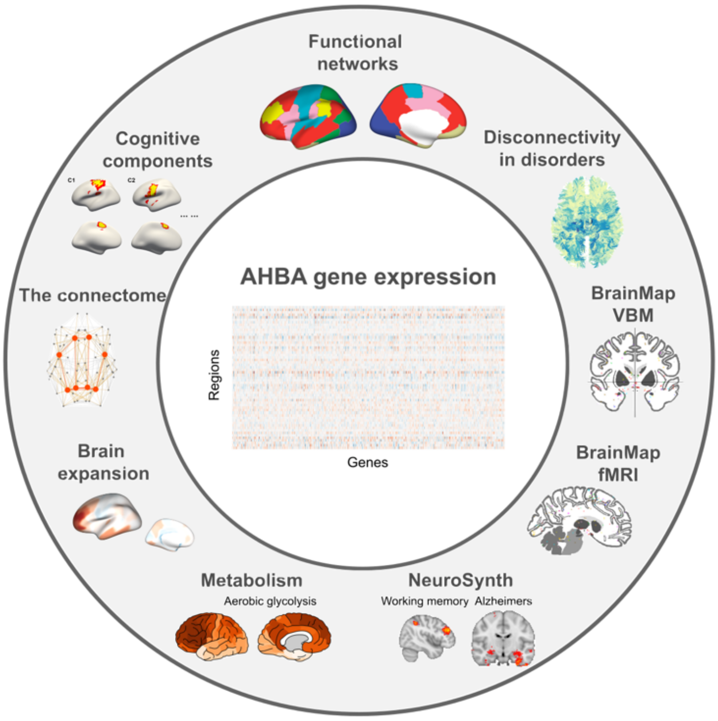


Figure S3. An overview of the preset imaging-derived phenotypes included in GAMBA. Brain gene expressions are extracted from the Allen Human Brain Atlas (AHBA) and are cross-referenced to distinct imaging-derived brain phenotypes. Brain phenotypes include (counterclockwise) functional networks, cognitive components, metrics of the connectome, evolutionary brain expression, and metabolic measurements in the healthy, and disconnectivity, structural and functional alterations in the diseased brain, as well as terms in NeuroSynth.

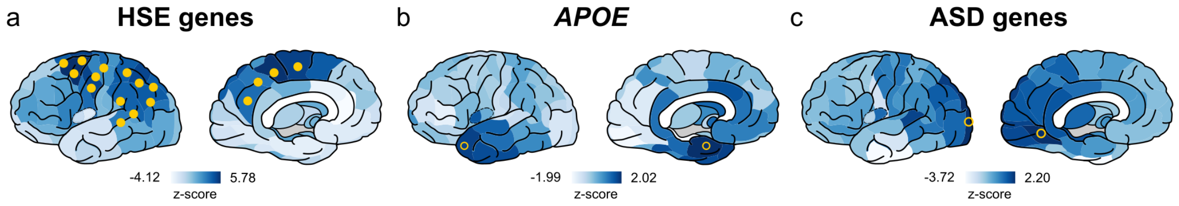


Figure S4. Brain maps of *z*-scores showing to what extent the expression level of the GOI is higher than genes on average. (a) Significantly higher expressions of HSE genes than all genes on average within the inferior/superior parietal cortex, supramarginal cortex, precuneus, superior/middle frontal cortex, precentral gyrus, and paracentral cortex (FDR corrected across 64 brain regions). (b) No significant over-expression for *APOE*. (c) No significant over-expression for 24 ASD *genes*. Yellow dots indicate significant ($q < 0.05$, FDR corrected). Yellow circles indicate $p < 0.05$ (not corrected).
